# Elevation and topography shape the diversity of mesoscale landscape-level woody plant species found in temperate evergreen forests

**DOI:** 10.1101/2023.03.29.534815

**Authors:** Shuntaro Watanabe, Yuri Maesako, Tomoya Inada

## Abstract

Plant species richness is influenced by complex interactions between biotic and abiotic factors that operate on different spatial scales. Since spatial scales vary continuously in nature, it is expected that multiple factors simultaneously affect species richness and composition at an intermediate spatial scale (i.e., the mesoscale landscape level). Previous studies have shown that local topography and elevation are important factors for shaping mesoscale landscape-level plant species richness; however, the relative importance of these factors has rarely been examined. Here, we used spatially explicit woody plant survey data to investigate the relative importance of topography, elevation, and disturbance at the mesoscale landscape level. We found that topography and elevation are important drivers of plant species richness and composition at the mesoscale landscape level and affect different components (i.e., the number of species and species composition, respectively). Our study also found that closely-related species coexisted along the elevational gradient, suggesting that niche partitioning among closely-related species is a fundamentally important feature of mesoscale species richness pattern. Furthermore, we found that specialization in a habitat of closely-related species can be established even within a limited environmental gradient. This suggests that biotic interactions among closely-related species may be an important factor driving habitat specialization, and biotic interactions may play an important role in shaping landscape-scale biodiversity patterns.

## Introduction

Plant species richness and composition are influenced by the complex interactions between biotic and abiotic factors. It is widely accepted that biotic and abiotic factors operate on different spatial scales. At the local or plot scale, patterns of plant diversity are influenced by biotic interactions (Gross et al., 2013) and/or disturbance regimes (Sipe and Bazzaz 1995). At larger spatial scales, habitat associations are strong and determine compositional shifts during changes in environmental conditions (Wiens & Donoghue 2004; Chase 2014). Since spatial scales vary continuously in nature, it is expected that biotic interactions and environmental factors simultaneously affect species richness and composition at intermediate spatial scales. Previous studies termed this intermediate spatial scale (1–100 km^2^) as the “mesoscale landscape” (Heikkinen 1996; Clark et al., 1996). Since mesoscale landscape forest processes link local gap dynamics to macroscale species distributions in a hierarchical manner, research at this scale is needed (Druckenbrod et al., 2019).

Numerous studies have revealed that local topography and elevation are important factors in shaping mesoscale landscape-level plant species richness. Since topographic features correlate with variables that are more directly related to plant resources, they are regarded as a good predictor of habitat. For example, topographic features such as slope and aspect often correlate with the distribution of nutrients (John et al. 2007) and soil moisture (Daws et al. 2002). Additionally, topographic features often correlated with disturbance regimes. Accordingly, species diversity and composition change along topographic gradients, from ridge to valley and from hilltop to hollow (e.g., Itoh et al. 1997; Enoki 2003, but see Suzuki 2011).

Changes in species richness along elevation gradients have also been documented in numerous studies (e.g. Rahbek, 1995; Grytnes and Vetaas, 2002; Aiba et al., 2004). Macroecological studies have shown that orographically enhanced precipitation, elevational and climatic gradients, and environmental heterogeneity are the key features linking high biodiversity to elevation (Antonelli et al., 2018). Previous studies have also reported that plant species richness is positively skewed (hump shaped) or monotonically decreases along elevation gradients (e.g. Aiba et al., 2001; Bruun et al., 2006; Rahbek, 2005). Furthermore, the determinant factors of community composition and diversity differ across distinct forest strata (e.g. overstorey, understorey; Bruun et al., 2006; Luo et al., 2016). Unlike topographic gradients, elevational gradients often create biological patterns that are mediated by interspecific interactions (e.g. competition; McCain and Grytnes 2010). These findings suggest that plant species diversity in heterogeneous forests is characterized by local or microscale environmental gradients that are mediated by elevational and topographic configurations (Tsujino et al. 2006).

Despite the studies mentioned above, relatively few studies have examined the relative importance of local topography and elevation at the mesoscale landscape level. Dearborn and Danby (2017) showed that plant communities in ecotone tundra varied more significantly with slope aspect than elevation. However, some studies have shown that both elevation and local topography affect plant richness (e.g. Tsujino and Yumoto 2013; Irl et al., 2015). Additionally, the relative importance of these factors could differ among plant life-form groups (Tsujino and Yumoto 2013; Luo et al., 2016).

In this context, we aimed to assess how topographic and elevational variables individually contribute to plant species richness at the mesoscale landscape level. We comprehensively investigated the diversity of mesoscale landscape-level (1 km^2^) plant species in the Kasugayama primary forest, which is a primary warm-temperate evergreen forest located in central Japan. The Kasugayama primary forest encompasses diverse micro-habitats attributed to the topographical variation and elevational variation and displays variation in canopy gap size distribution. We addressed the following questions: (1) What factors dictate tree species diversity at the mesoscale landscape level? (2) Is the number of species or species composition influenced by the same or different factors? (3) Do species behave as generalists, that is, do they occur throughout the environmental gradient within the study site, or do biotic and abiotic environments favor niche differentiation among species?

## Material and Method

### Study area

The Kasugayama primary forest has a total area of 250 ha and is located in Nara City, western Japan (34° 41’ N, 135° 51’ E). The elevation in the region ranges from 100 to 496 m. Since the forest has been preserved as a holy site of the Kasuga Taisha Shrine, hunting and logging have been prohibited since 841 AD (Maesako et al. 2007). In 2019, the mean annual temperature was 16.3 °C, and the average annual precipitation was 1482.5 mm. The highest point in the forest is 498 m. The natural vegetation in the area is evergreen broadleaved forests (Naka 1982); however, the deer population has recently increased in the forest, causing the spread of alien species, such as *Sapium sebiferum* and *Nagia nagi* (Maesako et al. 2007).

### Field survey

Field studies were conducted from June to September 2015. In the study area, 30 transect plots (∼0.1 ha in size) were established within range of 1 km2 (Fig. 1). Tree species richness was surveyed in each plot; all tree species with heights >130 cm and numbers of individuals were recorded. We measured the stem girth at breast height (1.3 m above ground) and calculated the diameter at breast height(DBH)from the girth. Species were grouped into three life-form categories, namely trees, shrubs, and climbing plants, according to Satake (1981–1982).

**Fig. 1.**
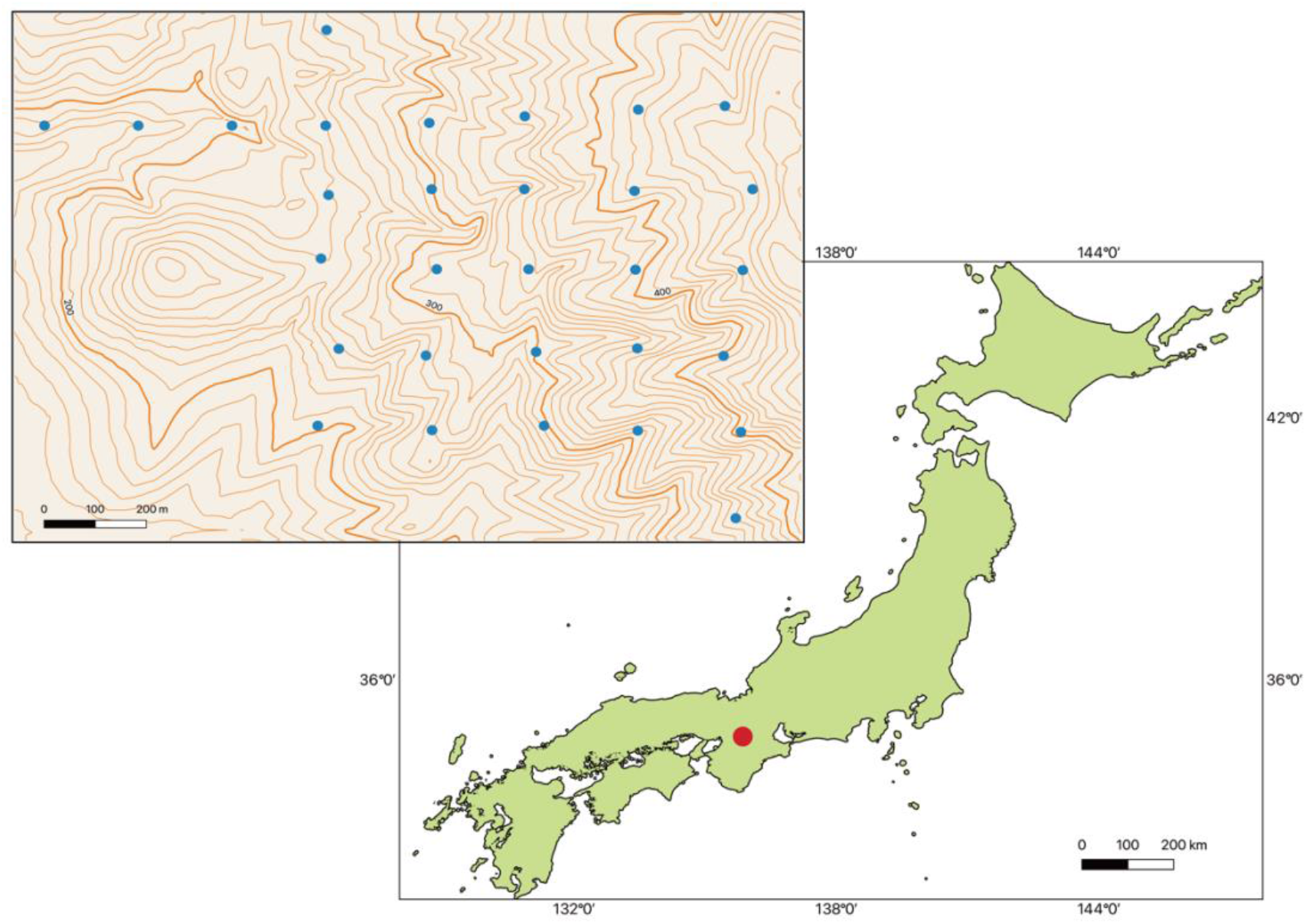
Location of the study site (∼1 km^2^) at the Kasugayama primary forest, Nara prefecture, Western Japan. The specific locations of the study plots are denoted by dots. Scale bar: 200 m.

### Environmental variables

Environmental variables were collected from each transect plot. Elevations were obtained using GPS. The slope inclination, Laplacian (i.e., an index of concavity and convexity of the ground), and slope aspect were obtained from a 20 m x 20 m digital elevation model using QGIS. Tangent transformation was applied to slope inclination. Sine and cosine transformations are applied to the slope aspect. In the sine transform, the values range from − 1 (east-facing) to 1 (west-facing), whereas in the cosine transform, values range from −1 (north-facing) to 1 (south-facing).

Canopy openness (%) at the plot scale was determined using hemispherical photographs and used as a proxy for recent disturbance. Hemispherical photographs were obtained using a Coolpix 8400 digital camera (Nikon, Tokyo, Japan) with an FC-E9 0.2x fisheye converter lens (Nikon, Tokyo, Japan). The camera was mounted on a tripod and oriented so that the top of each photograph faced the magnetic north. The lens was positioned at a height of 1.2 m, and photographs were taken using the open-sky reference method (Tani et al., 2011; Inada et al., 2017). We analyzed the color hemispherical photos using the Gap Light Analyzer software (ver. 2.02; Frazer et al., 1999).

### Statistical analysis

We used a generalized linear model (GLM) to assess the relationships between the number of species and environmental variables in the transect plots. A Poisson distribution was assumed for the response variables, and a generalized linear regression was fitted. For each response variable, after constructing models for all combinations of environmental variables, we identified the model with the smallest Akaike information criterion (AIC) as the best model and detected the other top three models.

To categorize the species distribution pattern across the elevational gradient, we used a generalized linear model that assumed a binomial distribution. We used the presence/absence of each species in each transect plot as a response variable. We categorized the distribution patterns of each species into four categories: (1) High-elevation species (species whose presence/absence patterns are positively correlated with elevation), (2) low-elevation species (species whose presence/absence patterns are negatively correlated with elevation), (3) generalist species (species whose presence/absence patterns are not significantly correlated with elevation), and (4) rare species (species that present less than three plots).

To identify differences in plant community composition among transect plots, we used non-metric multidimensional scaling (NMDS) ordination. We performed using the vegan library (Oksanen et al. 2020) in R (version 4.1.2). To determine whether elevation, topographic factors, and canopy openness were associated with differences in overall plant community composition, we ran a PerMANOVA on the results of each ordination with abiotic predictor variables. We used Vegan’s envfit function, which evaluates multidimensional correlations between NMDS site scores and independent environmental predictors, to further evaluate and visualize the effects of environmental variables on plant community composition.

## Results

### Factors affecting number of tree species

We recorded 70 species at the study site, including 49 tree species, 13 shrub species, and eight climbing plant species (S1). *Castanopsis cuspidata* was the most abundant species according to total basal area and *Quercus sessilifolia, Abies firma* and *Cryptomeria japonica* were also abundant (S1). *Neolitsea aciculate* and *Cleyera japonica* were the most abundant species according to their number of stems (S1).

In the best models for the number of woody plant species, the slope aspect was selected (Table 1). Slope aspect was selected in the models for tree and climbing plant species with lower AIC values, and it was positively correlated with the number of species. Accordingly, the number of species tended to be higher in the plots with steep slopes. Additionally, the total number of species and number of tree species were best explained by slope steepness alone, suggesting its primary importance.

**Table 1.**
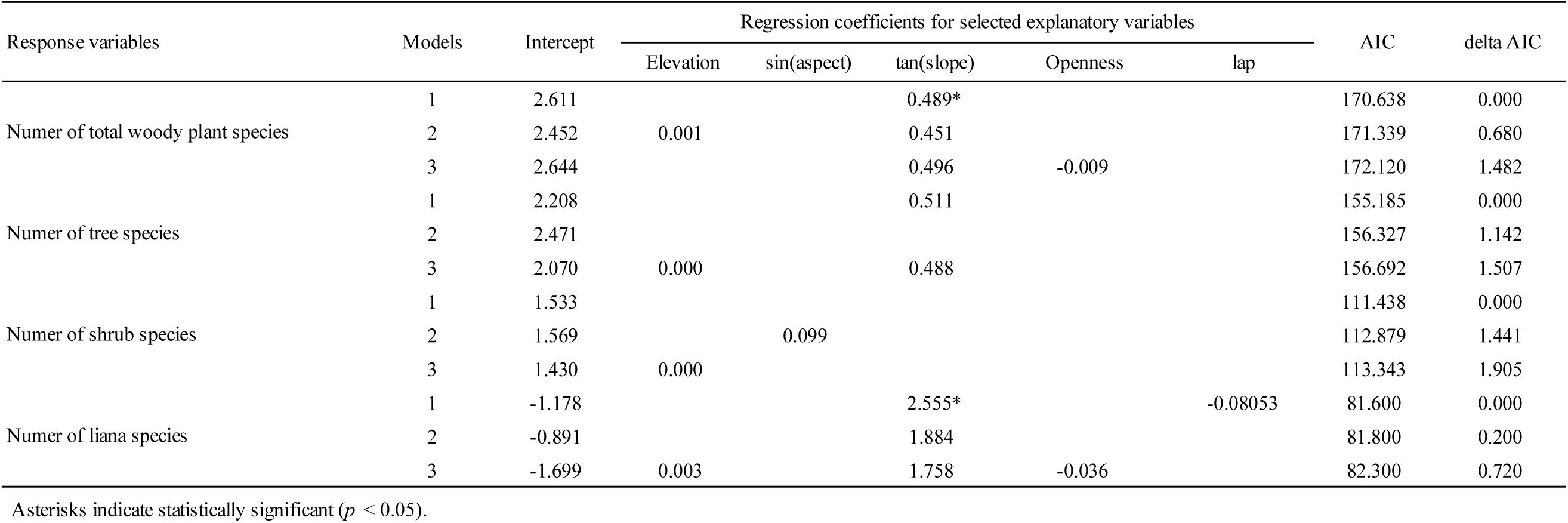
Top three significant generalized linear models for number of species in subpolots. AIC, Akaike information criterion; tan(slope), tangent transformed slope inclination; sin(aspect), sign transformed slope aspect; Openness, relative openness; lap, convex/concavity.

### Factors affect species composition and distribution

In the NMDS analysis for species composition, elevation showed significant relationships with the similarity in woody plant species composition among the transect plots (Fig. 2).

**Fig. 2.**
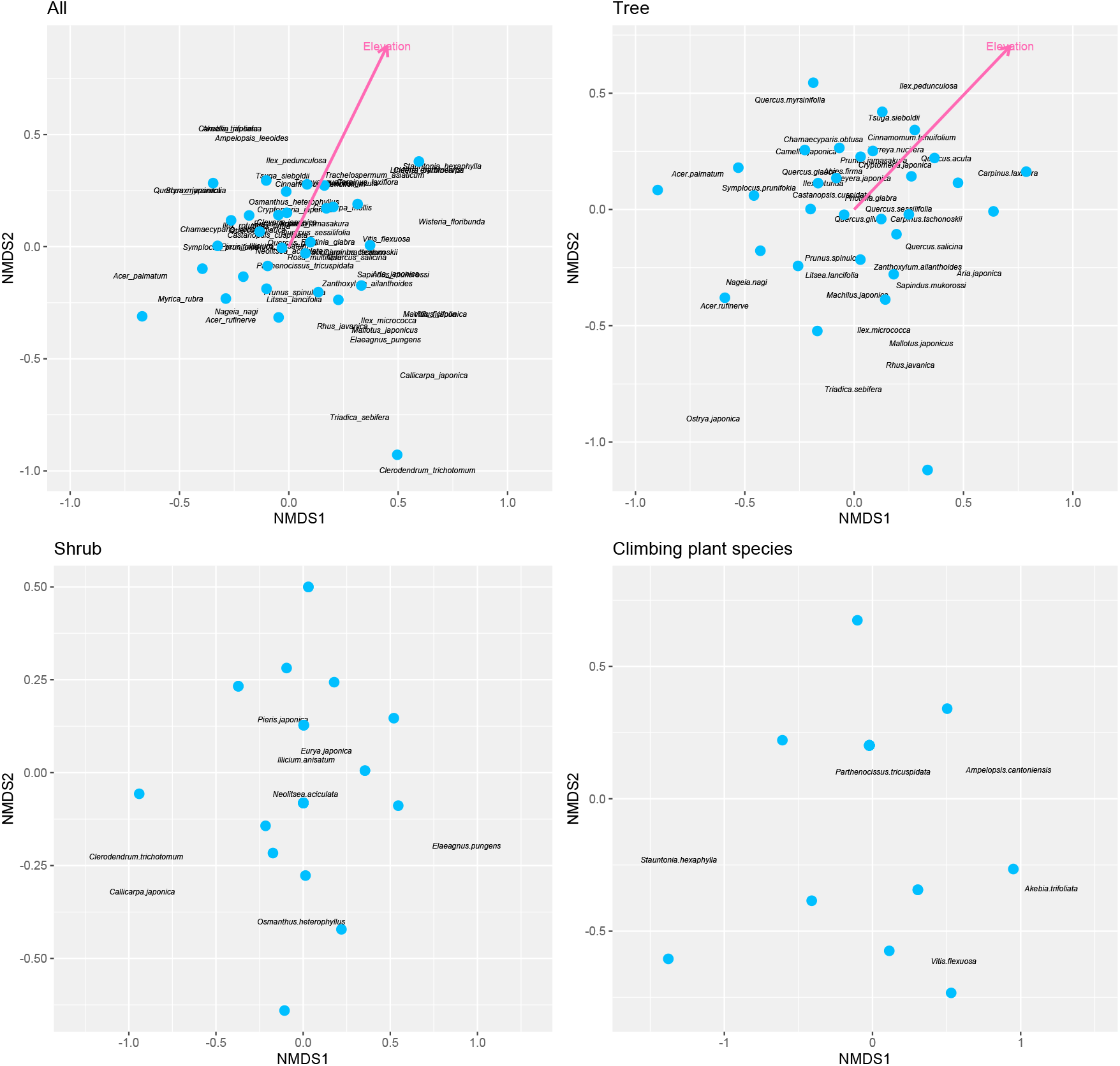
Differences in species composition of woody plant species determined using a non-metric multidimensional scaling (NMDS) and permutational multivariate analysis of variance (perMANOVA).

In the logistic regression analysis for species distribution, we detected 7 high-altitude species, 1 low-altitude species, 32 generalist species, and 36 rare species (S2). Three of the seven species categorized as high-elevation species had another species of the same genus categorized as generalists. For example, in *Quercus*, three of the six species were categorized as generalists and two were categorized as high-altitude species (Fig. 3). This suggests that niche partitioning with elevation occurs among congeneric species.

**Fig. 3.**
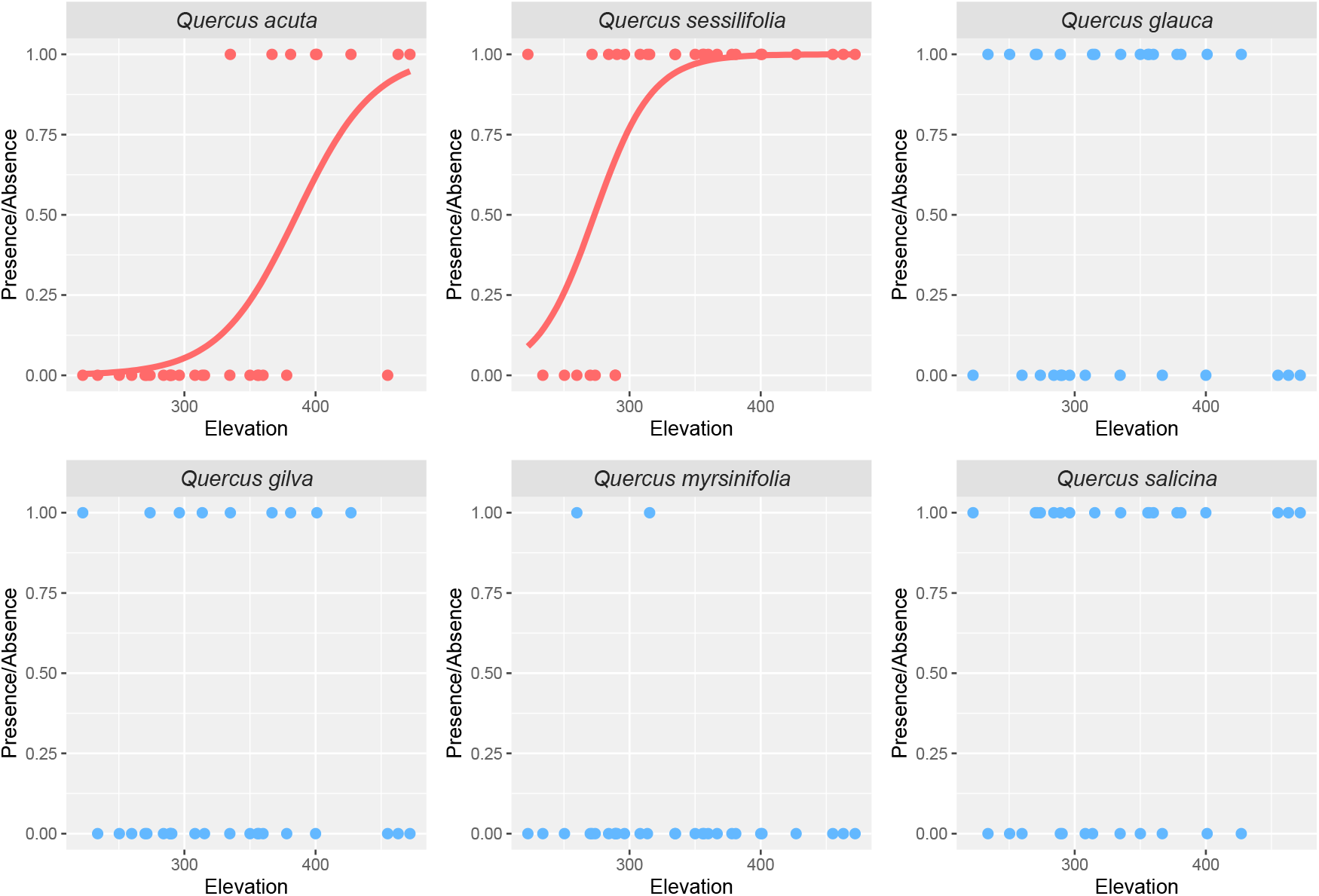
Distribution patterns of the Quercus along elevations in the study area. Curves indicate estimates of generalized linear model (*p* < 0.05).

## Discussion

### Effects of micro topography on species diversity

Our results suggest that slope steepness has a significant effect on tree species richness (Table 1). The effect of slope steepness on species richness differed among plant life forms, and the number of species tended to increase as the slope became steeper, particularly for climbing plant species. Previous studies have proposed two major mechanisms by which microtopography controls species diversity. One is that differences in soil properties along a topographic gradient affect plant diversity (e.g. Jones et al. 2008, Qin et al. 2019). Another idea is that topographic changes can modify species diversity by causing physical disturbances through landslides (e.g. Enoki 1998).

In this study, we found that tree species richness tended to be higher at steeper slope sites, especially for climbing plants. The frequency of disturbances such as landslides increases in areas with concave slopes (Hunter and Parker 1993). Ledo and Schmezer (2014) reported that population density and species diversity of tropical lianas are mainly determined by the intensity and frequency of disturbance. They reported that lianas maintain species diversity against disturbance by clonally increasing and becoming spatially coherent in response to disturbance. In this study, we also found a trend, although not statistically significant, for the climbing plant stem density to increase on steep slopes(S3), which is consistent with the results of Ledo and Schmezer (2014). These findings suggest that the patterns observed in this study might be driven by differences in disturbance frequency owing to differences in microtopography.

Previous studies have pointed out that canopy gaps often facilitate the establishment and growth of many tree species (Schnitzer and Carson 2001). However, we did not find a consistent relationship between canopy openness and species diversity (Table 2). One possible reason is the influence of deer grazing. Deer population density has been increasing in this study area since the 1990s. In our study area, the deer abundance was significantly increased around tree fall gap and deer grazing strongly regulate the regeneration of pioneer tree species (Shimoda et al., 1994; Maesako et al., 2018). These results suggest that gap dynamics, which have been widely regarded as major mechanisms for the maintenance of species diversity, do not work well in our study area. Additionally, it is reported that the deer population density is often affected by topography, and deer population density tends to a decrease in steep slope area (Ganskopp and Vavra 1987). The increase in species richness at steeper slope sites in our study possibly reflects the heterogeneity of deer population density and herbivory intensity.

### Factors affecting species composition

The results of the NMDS analysis suggested that tree species composition at this study site changed with elevation. Additionally, the results of the regression analysis assuming a binomial distribution revealed that habitat partitioning along the elevational gradient was frequently observed among congeneric species (Fig.3, S2). These findings suggest that habitat specialization, especially among closely-related species, is an important driver of changes in species composition with elevation.

Previous studies have reported that water-use strategies, edaphic conditions, and light conditions often shape divergent habitat associations among congeneric species pairs (Itoh et al., 1998, Baltzer et al. 2005, Yamasaki et al., 2018). Schmitt et al. (2021) reported that the coexistence of related species in tropical forests is explained by both species-specific habitat specialization within species complexes and the broad ecological distribution of species complexes, and species more tolerant to competition for resources grow in drier and less fertile plateaus and slopes. On the other hand, Christie and Strauss (2020) evaluated the factor that involves niche-partition among closely-related species using a transplant experiment and found that micro-parapatry, which inhabits continuous or adjacent habitat patches and occurs within the seed dispersal range, yet rarely overlaps in fine-scale distribution, of two closely-related and reproductively isolated *Streptanthus* species is determined by negative reproductive interactions among species.

The distribution pattern along the elevation for each species in our study area can be classified into two major types: generalist and high-elevation. Interestingly, both the generalist and high-elevation types were included in the same genus (Fig. 3, S2). This suggests that lower elevation sites tend to be dominated by a particular species, whereas multiple closely-related species tend to coexist at higher elevation sites. Elevation generally corresponds to temperature, water availability, and/or edaphic conditions, and higher elevation sites are generally more unsuitable for plants than low-altitude areas. A recent meta-analysis highlighted that the cool range limit (range limit toward higher elevation) is often constrained by abiotic factors, whereas the warm range limit (range limit toward lower altitude) is often determined by biotic factors (Paquette and Hargreves 2021). Among the biotic interactions, negative interspecific interactions, such as competition or reproductive interference, are known to hamper the coexistence of species at a fine scale (e.g. Silvertown 2004; Takakura et al., 2009; Christie and Strauss 2020). Theoretical studies assuming host specialization of herbivorous insects suggest that specialization in habitats is most likely to occur when competition and negative reproductive interactions among closely-related species are at intermediate levels (Nishida et al., 2016). These facts suggest that negative interspecific interactions possibly involve habitat specialization among congeneric tree species, and that elevational gradients affect species composition and distribution at the mesoscale landscape level through interspecific interactions. Watanabe and Maesako (2021) analyzed the distribution of *Quercus* and *Carpinus* in the study area and suggested that the effects of negative interactions among congeneric tree species were possibly weaker than those reported in previous studies. The mechanism by which the effects of negative interactions among congeneric species are not well understood, but it is quite likely that the mitigation of negative interactions among congeneric species is responsible for specialization of particular species into unsuitable environments (high altitudes) at the mesoscale landscape level. Further theoretical and empirical research is required to determine how interspecific interactions are related to niche partitioning at the mesoscale.

## Conclusion

We found that topography and elevation are important drivers of plant species richness and composition at the mesoscale landscape level, and they affect different components, namely the number of species and species composition, respectively. Our study also found that closely-related species coexisted along the elevational gradient, suggesting that niche partitioning among closely-related species is a fundamentally important feature of mesoscale species richness patterns. Furthermore, we found that specialization in the habitat of closely-related species can be established even within a limited environmental gradient. This suggests that biotic interactions among closely-related species may be an important factor driving habitat specialization, and that biotic interactions may play an important role in shaping landscape-scale biodiversity patterns.

## Acknowledgement

We would like to thank Minori Hikichi and Kayo Takasu for fieldwork assistance. This work was supported by JSPS KAKENHI (grant number JP 23510300).

## Supplemental material

S1 Life form, basal are and stem numbers of woody plant in this study site.

S2 Distribution pattern of woody plant in this study site.

S3 Relationship between slope steepness and stem density.

## Notes

### Competing Interest Statement

The authors have declared no competing interest.

